# Locally-adapted *Mimulus* ecotypes differentially impact rhizosphere bacterial and archaeal communities in an environment-dependent manner

**DOI:** 10.1101/652883

**Authors:** Alan W. Bowsher, Patrick J. Kearns, Damian Popovic, David B. Lowry, Ashley Shade

**Affiliations:** Department of Microbiology and Molecular Genetics, Michigan State University, East Lansing, MI; Plant Resilience Institute, Michigan State University, East Lansing, MI; Department of Plant Biology, Michigan State University, East Lansing, MI; Program in Ecology, Evolutionary Biology and Behavior, Michigan State University, East Lansing, MI; DOE Great Lakes Bioenergy Research Center, Michigan State University, East Lansing, MI

**Author notes:** Corresponding author: A. Shade.

**Keywords:** root microbiome, *Mimulus guttatus*, 16S rRNA gene, plant-microbe interactions

## Abstract

Plant root-microbe interactions influence plant productivity, health, and resistance to stress. Although there is evidence that plant species and even genotypes can alter soil microbial community structure, environmental conditions can potentially outweigh plant genetic effects. Here, we used a reciprocal transplant experiment to understand the contributions of the environment and the host plant to rhizosphere microbiome composition in locally-adapted ecotypes of *Mimulus guttatus* (syn. *Erythranthe guttata* (Fisch. ex DC.) G.L. Nesom). Two genotypes of a coastal ecotype and two genotypes of an inland ecotype were planted at coastal and inland sites. After three months, we collected rhizosphere and bulk soil and assessed microbial communities by 16S rRNA gene sequencing. We found that local environment (coastal versus inland site) strongly influenced rhizosphere communities, at least in part due to distinct local microbial species pools. Host identity played a smaller role: at each site, the ecotypes exhibited remarkably similar composition of microbial communities at the class level, indicating that divergent *M. guttatus* ecotypes recruit phylogenetically similar rhizosphere communities, even in environments to which they are maladapted. Nevertheless, the two ecotypes significantly differed in community composition at the inland site due to an exclusive set of rare taxa associated with each ecotype. Although our results indicate that locally-adapted *M. guttatus* ecotypes are genetically diverged in factors shaping rhizosphere communities, environmental factors can trump genetic factors in shaping the *M. guttatus* microbiome. Overall, our findings demonstrate that wild plants strongly impact root-associated microbial communities, but hierarchical drivers interact to shape microbial community assembly outcomes.

## Introduction

The rhizosphere (the narrow zone of soil surrounding plant roots) is a highly diverse and active microenvironment. In addition to influencing soil structure, moisture, and nutrient availability (Marschner et al. 1987; Angers and Caron 1998; McKinney and Cleland 2014), plant roots continuously supply labile carbon to the soil through root exudation. These continual carbon inputs recruit a host of soil microbes to the rhizosphere (Bressan et al. 2009; Bulgarelli et al. 2012; Chaparro et al. 2014; Zhalnina et al. 2018), often resulting in distinct microbial communities compared to the surrounding bulk soil (Berendsen et al. 2012; Bever et al. 2012; Philippot et al. 2013). Rhizosphere microbial communities can strongly impact plant health and productivity, altering plant morphology (Friesen et al. 2011), phenology (Wagner et al. 2014), and plant resistance to both biotic (Santhanam et al. 2015; Busby et al. 2016; Ritpitakphong et al. 2016) and abiotic stresses (Lau and Lennon 2011, 2012). Nevertheless, despite the critical importance of rhizosphere communities for plant productivity, the factors shaping the rhizosphere microbiome are complex and not fully understood (Berg and Smalla 2009; Lareen et al. 2016; Sasse et al. 2018).

One factor that can strongly influence rhizosphere community composition is plant host identity. Plant species and even genotypes within species can differ in rhizosphere community structure when planted in a common environment (Aira et al. 2010; Bouffaud et al. 2012; Edwards et al. 2015; Mahoney et al. 2017; Berg et al. 2002; Bowen et al. 2017; Fitzpatrick et al. 2018). This finding is often suggested to result, at least in part, from species-specific root exudation patterns recruiting different community members (Marschner et al. 2001). Indeed, numerous studies suggest root exudation is the primary mechanism by which plants mediate rhizosphere community assembly and function (Broeckling et al. 2008; Haichar et al. 2008; Carvalhais et al. 2015; Hu et al. 2018). Other species- or genotype-specific factors could also contribute, such as differences in rooting depth (Aleklett et al. 2015) and root architecture (Pérez-Jaramillo et al. 2017), given that microbial community composition can shift with soil depth (Fierer et al. 2003; Ko et al. 2017).

In addition to the influence of plant host identity, environmental factors can also shape the rhizosphere microbiome. For example, the local environment directly affects rhizosphere communities by determining the available source pool of microorganisms, since soil microbial communities are structured by both spatial and environmental gradients (Fierer and Jackson 2006; Xue et al. 2018; Rath et al. 2019). Local environmental conditions can also indirectly influence rhizosphere community composition by affecting plant and microbial physiology (Aira et al. 2010). For example, many environmental factors can influence root exudate composition, such as nutrient availability (Zhang et al. 1991; Carvalhais et al. 2011), pathogenesis (Gu et al. 2016), drought (Gargallo-Garriga et al. 2018), and flooding (Henry et al. 2007), thereby influencing rhizosphere composition. As a result, environmental conditions can outweigh the effects of plant host identity (i.e. differences among plant species or genotypes) in structuring rhizosphere communities (Marschner et al. 2004; Peiffer et al. 2013). While considerable recent microbiome research has been focused on economically important crops, less is known about the interplay between plant host and the local environment for wild plants, which experience relatively higher variability in their local environments than plants grown in managed systems.

In this study, we used a field reciprocal transplant experiment to better understand the contributions of both the environment and host plant identity to rhizosphere microbiome composition. We used two locally adapted ecotypes (coastal versus inland) of the yellow monkeyflower, *Mimulus guttatu*s (syn. *Erythranthe guttata* (Fisch. ex DC.) G.L. Nesom), a model species for ecological and evolutionary genomics (Twyford et al. 2015; Wu et al. 2008). Coastal and inland ecotypes are highly locally adapted to their respective habitats (Hall et al. 2010; Lowry et al. 2008; Lowry and Willis 2010; Hall and Willis 2006). Inland habitats of *M. guttatus* experience a hot summer drought, for which these populations have evolved an early flowering, annual life-history strategy to escape from the long period of low soil water availability (Lowry et al. 2008; Hall and Willis 2006). In contrast, coastal habitats typically are much cooler as a result of proximity to the Pacific Ocean, which drives the production of summer sea fog. However, coastal populations of *M. guttatus* contend with pervasive oceanic salt spray, for which they are locally adapted (Lowry et al. 2008, 2009). Here, we planted coastal and inland ecotypes of *M. guttatus* in both coastal and inland sites and investigated rhizosphere and bulk soil microbial community composition after three months of growth.

## Materials and Methods

### Experimental Design

To establish the relative role of environment (coastal versus inland site) and ecotype (coastal perennial versus inland annual) on the *M. guttatus* microbial rhizosphere community, we leveraged a reciprocal transplant experiment conducted in Sonoma County, CA, USA in the spring of 2017 (Popovic and Lowry 2019). Briefly, accessions from two coastal perennial populations (SWB-11, 39.0359 N, - 123.6905 W; MRR-13, 38.4564 N, −123.1409 W) and two inland annual populations (LMC-24, 38.8640 N, −123.0840 W; OCC-31, 38.4095 N, −122.9355 W) were used for the experiment. Source populations for the SWB and LMC seeds are in Mendocino County, CA, and have been used in many recent studies of genetics and local adaptation in this system (Lowry et al. 2008, 2009). The MRR and OCC source populations are located in Sonoma County, CA (Popovic and Lowry 2019). All accessions were grown for at least one generation in the Michigan State University greenhouses to control for maternal effects.

Seeds were planted on wet Sunshine Soil Mix #1 (SunGro Horticulture, Agawam, MA) on February 1, 2017 in two 54.28 × 27.94 cm potting trays per genotype. Seeds were then stratified at 4°C for 10-17 days (10 days for coastal accessions, 17 days for inland accessions), and subsequently germinated at University of California, Berkeley’s Oxford Track greenhouse facilities under 16 hours of light. Different lengths of stratification were used for the two ecotypes because the inland ecotype germinates earlier and grows faster than the coastal genotype early in development. This allowed seedlings to be transplanted to the field later at the same developmental stage. On February 28th, all seedlings were moved to the greenhouse at the Bodega Marine Reserve (bml.ucdavis.edu/bmr/) in Bodega Bay, CA.

We transplanted seedlings at the four-leaf stage into the coastal site on March 8th and into the inland site on March 9th. The coastal site was located at the Bodega Marine Reserve, Bodega Bay, CA, in a perennial seep at the south end of Horseshoe Cove (38.315716 N, −123.068625 W; ∼60 m from the ocean). The inland site was planted in a seasonal grassland seep at the Pepperwood Preserve in Santa Rosa, CA (38.575545 N, −122.700851 W; 39.84 km from the ocean). Native populations of *M. guttatus* are located in both seeps. Prior to planting, three 1 × 1 m plots were cleared of native vegetation at each site. Each plot included a total of 100 plants *(N=*25 of each genotype), which were all equally spaced from one another throughout the plot (*N=*100 per plot, 300 per site, 600 total). Plants were then grown for three months until being harvested for rhizosphere community analyses.

### Sample collection and processing

On June 13^th^-15^th^, five replicate *M. guttatus* rhizosphere soils were collected from each genotype at each field environment from plants that were spatially distributed across all three plots. Rhizosphere soil was isolated by uprooting the plant with a trowel, discarding excess soils from around the roots, and shaking what soil remained attached to the root into a sterile Whirl-Pak bag. Rhizosphere soils were homogenized with an ethanol-sterilized metal spatula, aliquoted into cryovials, flash frozen in liquid nitrogen, and stored on ice. Above- and belowground tissue for each plant was stored in a paper bag and transported at ambient temperature to the lab at Michigan State University, washed with distilled water, and dried for 1 week at 60°C before measuring dry biomass. In addition, bulk soil cores (10 cm x 2 cm) were collected randomly across the three plots at each site, sieved, and homogenized in a sterile Whirl-Pak bag and stored on ice. Bulk soil samples were subsequently analyzed for phosphorus, potassium, calcium, magnesium, copper, percent organic matter, sodium, nitrate, ammonium, percent nitrogen, and sulfur at the Michigan State University Soil and Plant Nutrient Laboratory following their standard protocols (http://www.spnl.msu.edu/). Gravimetric soil water content was determined from the loss of mass in soils dried for one week at 60°C. We assessed significant differences in soil chemistry with t-tests in R 3.5.0 (R Core Team 2018). The homogeneity of variance assumption was assessed using both Bartlett’s and Levene’s tests (Levene 1960; Snedecor and Cochran 1989) in the ‘car’ package (Fox and Weisberg 2011) of R, and the Welch’s t-test was used when the homogeneity of variance assumption was not met.

### DNA Extraction and Sequencing

DNA was extracted from the five replicate rhizosphere soil samples of each *M. guttatus* genotype from each environment (n=40 samples; five replicates of each of four genotypes at each of two sites), as well as from ten bulk soil samples (five replicates from each of two sites). We used the MoBio PowerSoil Total DNA Isolation Kit (Carlsbad, CA, USA) following the manufacturer’s instructions. Extracted DNA was quantified fluorometrically with the Qubit (ThermoFisher, Waltham, MA, USA). DNA from each sample was diluted to < 10 ng µl^−1^ for paired-end amplicon sequencing using the dual-indexed primer pair 515F/806R (Kozich et al. 2013). Samples were prepared for sequencing by the Michigan State University Genomics Core (East Lansing, MI, USA) including PCR amplification and library preparation using the Illumina TruSeq Nano DNA Library Preparation Kit. Paired-end, 250bp reads were generated on an Illumina MiSeq and the Genomics Core provided standard Illumina quality control and sample demultiplexing.

### Sequence processing

The rhizosphere and bulk soil sequencing datasets were analyzed together. Paired-end reads were merged using USEARCH v10.0.240 (Edgar 2010) and primer-binding regions removed using cutadapt v1.18 (Martin 2011), then reads were quality-filtered, dereplicated, and clustered into zero-radius OTUs using the USEARCH v9.2.64/v10.0.240 and UNOISE pipeline (Edgar 2016). Taxonomy annotations were assigned in Qiime v1.9.0 (Caporaso et al. 2010) using UCLUST (Edgar 2010) against the SILVA rRNA database v123 (Quast et al. 2013) and were added to the .biom file using the biom-format package (McDonald et al. 2012). Sequences that were unassigned at the phylum level, along with those matching chloroplasts or mitochondria, were excluded from analyses. Representative sequences were aligned using MUSCLE 3.8.1 (Edgar 2004) and FastTree v2.1.10 (Price et al. 2009, 2010) was used to build a phylogenetic tree. Samples were rarefied to the minimum number of sequences observed per sample (22,354) for all subsequent analyses. We calculated species richness, Shannon diversity, and phylogenetic diversity in QIIME, as well as beta diversity using weighted UniFrac distance (Lozupone and Knight 2005) for Principal Coordinates Analysis (PCoA).

Statistical analyses were performed in R 3.5.0 (R Core Team 2018). We assessed the effects of abiotic (phosphorus, potassium, calcium, magnesium, copper, percent organic matter, sodium, nitrate, ammonium, percent nitrogen, and sulfur) parameters on microbial community composition by fitting variables to weighted UniFrac distance with the R package vegan v2.5-2 (Oksanen et al. 2018). We included parameters that had significant explanatory value (p < 0.1) for PCoA axis 1 or 2. Differences in community composition across categorical groups (rhizosphere versus bulk soil, inland versus coastal sites, inland versus coastal ecotypes at each site, etc.) were calculated with PERMANOVA (Anderson 2001). We also tested for differences in group dispersions with PERMDISP (Anderson 2006). For alpha diversity metrics (species richness, phylogenetic diversity, and Shannon diversity), we tested for differences between ecotypes at each site, and between each ecotype and bulk soil at each site, using t-tests. Next, we selected the twenty most abundant taxa at the class level and tested for differences in abundance of these taxa using t-tests with an FDR-adjusted p-value for multiple comparisons. Within each site, we compared inland versus coastal ecotypes, as well as each ecotype versus bulk soil. We also compared genotypes within ecotypes at each site.

Given that the coastal and inland ecotypes differed in community composition only at the inland site, we further explored the inland site alone to better understand the factors distinguishing the microbiomes of the two ecotypes. First, we conducted an indicator species analysis, which aims to determine which taxa are characteristic of a given treatment group, taking into account the abundances of a given taxon for each treatment group (specificity), as well as the proportion of samples in each treatment group in which that taxon occurs (fidelity) (De Cáceres and Legendre 2009; De Cáceres et al. 2010). We used the multipatt function (De Cáceres et al. 2010) in the R package indicspecies (De Cáceres and Legendre 2009). Next, we tested for ecotype differences in relative abundance of individual OTUs using t-tests with FDR-adjusted p-values for multiple comparisons. Finally, we generated Venn diagrams using the R packages gplots (Warnes et al. 2019) and VennDiagram (Chen 2018) to assess differences in the presence/absence of individual taxa between the two ecotypes. Data were visualized using a combination of the R packages ggplot2 v2.2.1 (Wickham 2009), reshape2 v.1.4.3 (Wickham 2007), and cowplot v0.9.2 (Wilke 2017). Package plyr v.1.8.4 (Wickham 2011) was used for data summaries.

### Data availability and computing workflows

Raw reads were submitted to the NCBI Sequence Read Archive under accession numbers PRJNA451377 (rhizosphere samples) and PRJNA526056 (bulk soil samples). All plant and environmental data, as well as computational workflows and custom scripts, are available on GitHub (https://github.com/ShadeLab/PAPER_MimulusRecipTransplant_Submitted).

## Results

### Soil characteristics and plant performance differ across sites

The coastal and inland sites had very different soil properties (Table 1). Nearly all measured abiotic parameters significantly differed between the coastal and inland sites, with the exception of pH, ammonium, nitrate, and percent nitrogen. Plants also performed differently in the coastal and inland sites. Plants grown in the coastal site tended to be larger in both shoot and root mass than those grown in the inland site (Figure S1), although this was only significant for genotype MRR (coastal ecotype).

### Site and ecotype influence microbial community composition

A principal coordinates analysis based on weighted UniFrac distances found that two axes captured nearly 60% of the variation in the amplicon sequencing dataset (45.8% variation explained for PC1 and 13.9% for PC2) (Figure 1). Numerous abiotic parameters had significantly explanatory value for PCoA axis 1, which largely distinguished the coastal and the inland sites. Coastal site samples were associated with greater moisture content, sodium, phosphorus, and sulfur, while inland site samples were associated with greater potassium, calcium, magnesium, and copper (Figure 1).

**Figure 1.**
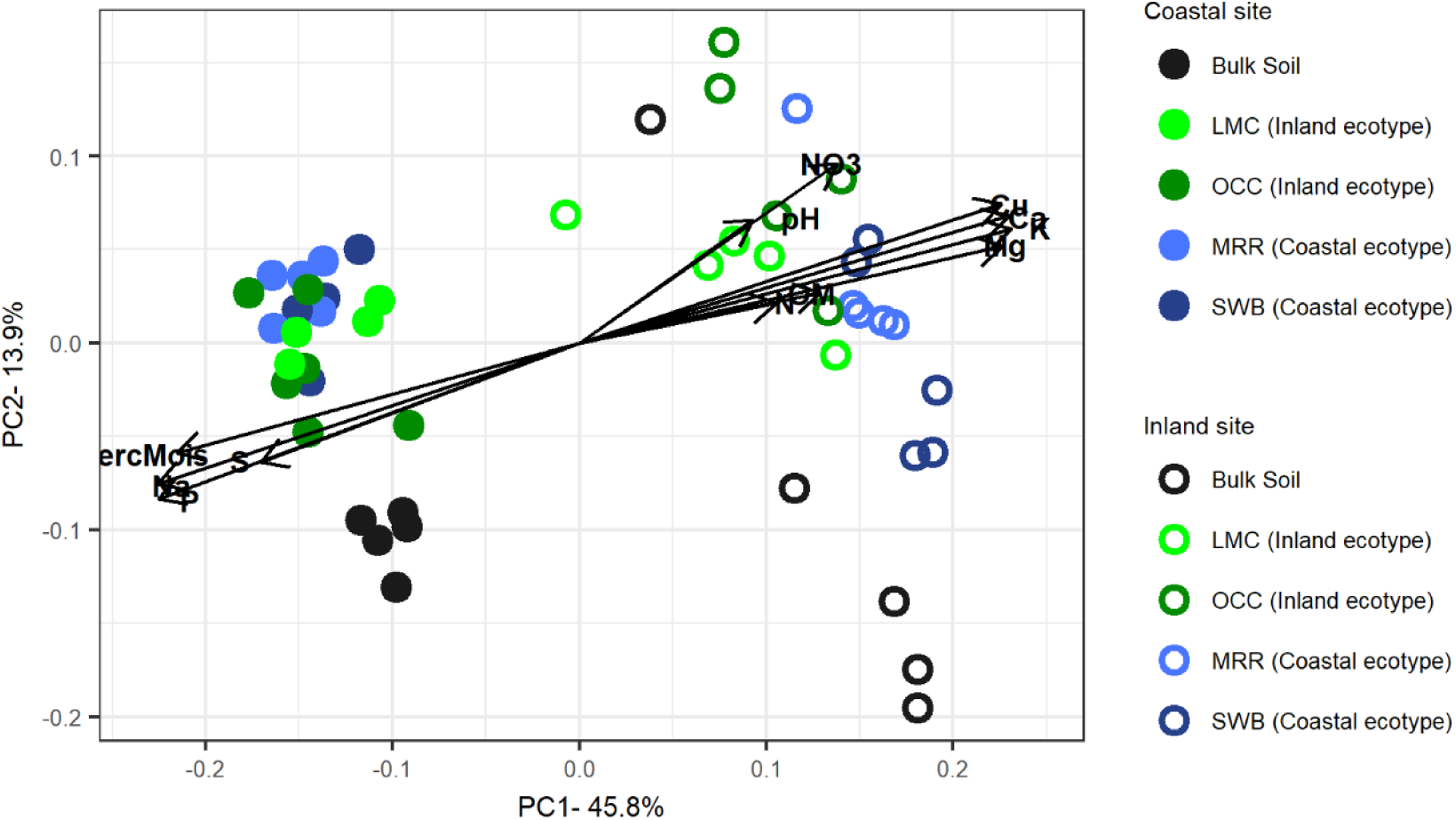
Principal coordinates analysis based on weighted UniFrac distances of bacterial and archaeal community structure. The strength of statistically significant (p < 0.05) explanatory variables are shown with solid arrows.

PERMANOVA revealed significant clustering of microbial communities by sample type (rhizosphere versus bulk soil; *F*=8.011, *P*=0.001) and site (coastal versus inland; *F*=43.227, *P* =0.001), as well as their interaction (*F*=4.307, *P* =0.006). We therefore investigated further by dividing the dataset by site and found that rhizosphere and bulk soils significantly differed in community composition at both the coastal and the inland sites (*F*=8.2951, *P*=0.001; and *F*=4.918, *P* =0.005, respectively), and differed in variability by PERMDISP at the coastal site (*F*=10.73, *P*=0.002). We next subdivided the rhizosphere samples by ecotype. We found that site influenced community composition for both the coastal (*F*=28.828, *P*=0.001) and inland ecotypes (*F*=16.319, *P*=0.001). In addition, the coastal ecotype differed in variability between the two sites (*F*=7.3244, *P*=0.013). Next, we found that inland ecotype rhizospheres differed from bulk soil in community composition at both the coastal and inland sites (*F*=6.2055, *P*=0.001; and *F*=5.2513, *P*=0.007, respectively), and differed in variability at the coastal site (*F*=13.198, *P*=0.004). Similarly, coastal ecotype rhizospheres differed from bulk soil at both the coastal and inland sites (*F*=10.474, *P*=0.001; and *F*=3.8461, *P*=0.004, respectively). We also tested for differences between ecotypes at each site and found that inland and coastal ecotypes differed in community composition at the inland site (*F*=3.279, *P*=0.006), but not at the coastal site (*F*=1.6859, *P*=0.095). Finally, we tested for differences between genotypes (within each ecotype at each site), and found that genotypes did not differ in any instance (all *P*>0.1).

### Ecotypes differ in rhizosphere communities at inland site

Across environments and ecotypes, we detected 14,869 OTUs spanning a breadth of phylogenetic diversity. Overall, alpha diversity metrics did not differ between either ecotype and bulk soil at either site (Figure 2). However, the inland ecotype exhibited greater species richness (*t*=-3.2507, *P*=0.006), phylogenetic diversity (*t*=-3.2446, *P*=0.004), and Shannon diversity (*t*=-2.9905, *P*=0.012) than the coastal ecotype at the inland site (Figure 2).

**Figure 2.**
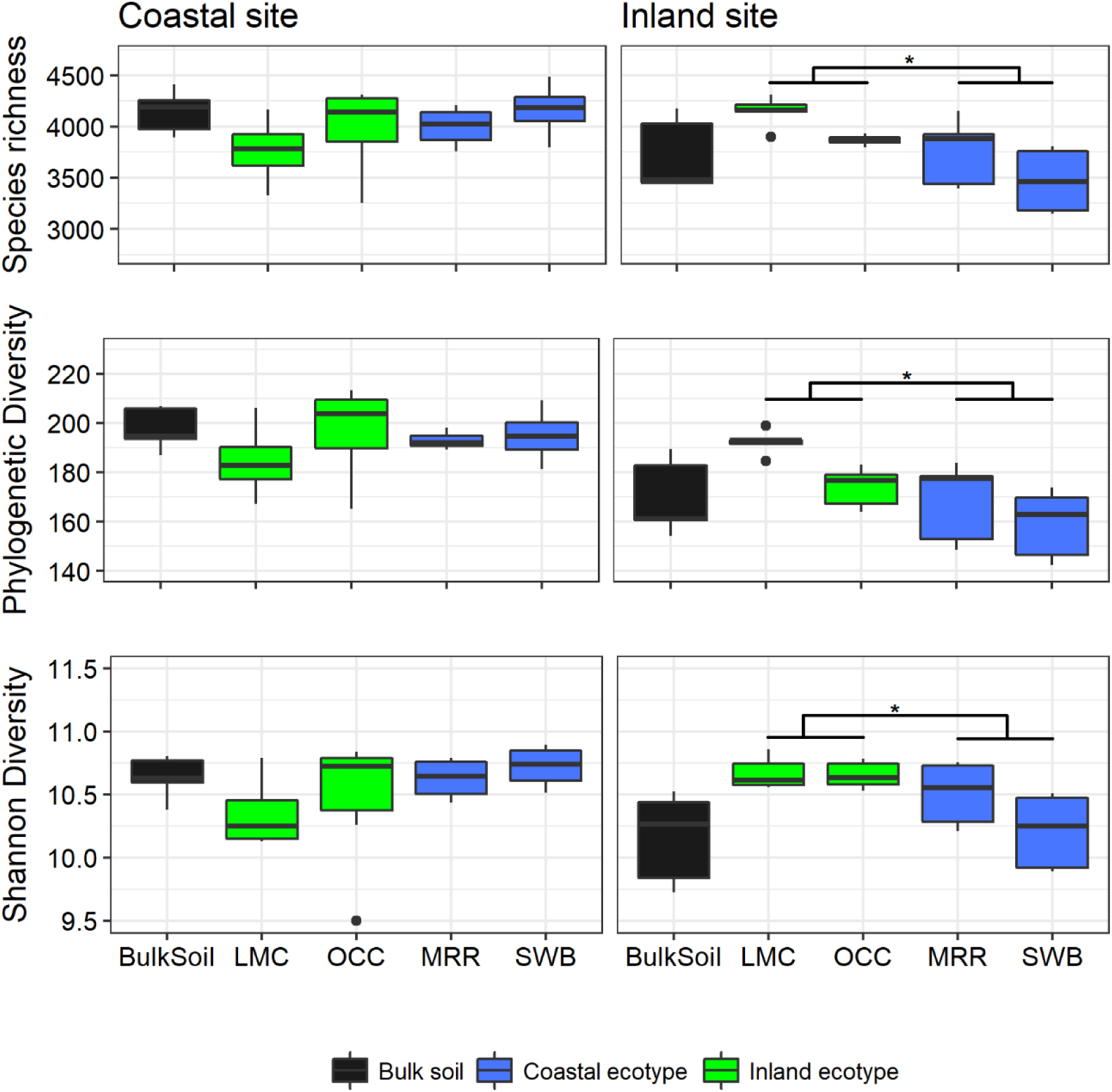
Metrics of alpha diversity in bulk soil and rhizosphere of coastal (genotypes MRR and SWB pooled) and inland (genotypes LMC and OCC pooled) ecotypes of *Mimulus guttatus* planted in two environments. Instances where ecotypes significantly differ are indicated with an asterisk (*).

Each ecotype exhibited some of the same compositional shifts in microbial communities (relative to bulk soil) in both sites. At both the coastal and inland sites, the inland ecotype exhibited lower relative abundance of Acidobacteria, Gemmatimonadetes, Nitrospira, and higher relative abundance of Planctomycetacia, compared to bulk soils (Figure 3). Similarly, at both sites, the coastal ecotype exhibited lower relative abundance of Nitrospira, and higher relative abundance of Planctomycetacia, compared to bulk soils. Within each site, both ecotypes influenced the relative abundance of numerous taxa in similar ways. At the coastal site, both ecotypes exhibited lower relative abundance of Acidobacteria, Anaerolineae, Gemmatimodetes, Nitrospira, Deltaproteobacteria, and OPB35-Soil, and higher relative abundance of Thermoleophilia, Cytophagia, Sphingobacteria, KD4-96, Planctomycetacia, and Alpha-proteobacteria, compared to bulk soil (Figure 3). Similarly, at inland site, both ecotypes exhibited lower relative abundance of Nitrospira and higher relative abundance of Planctomycetacia compared to bulk soil. There were exceptions to this rule, however. For example, at the inland site, the inland ecotype exhibited lower relative abundance of Acidobacteria, Gemmatimomdetes, Spartobacteria, and higher relative abundance of Actinobacteria compared to bulk soil, while the coastal ecotype did not (Figure 3).

**Figure 3.**
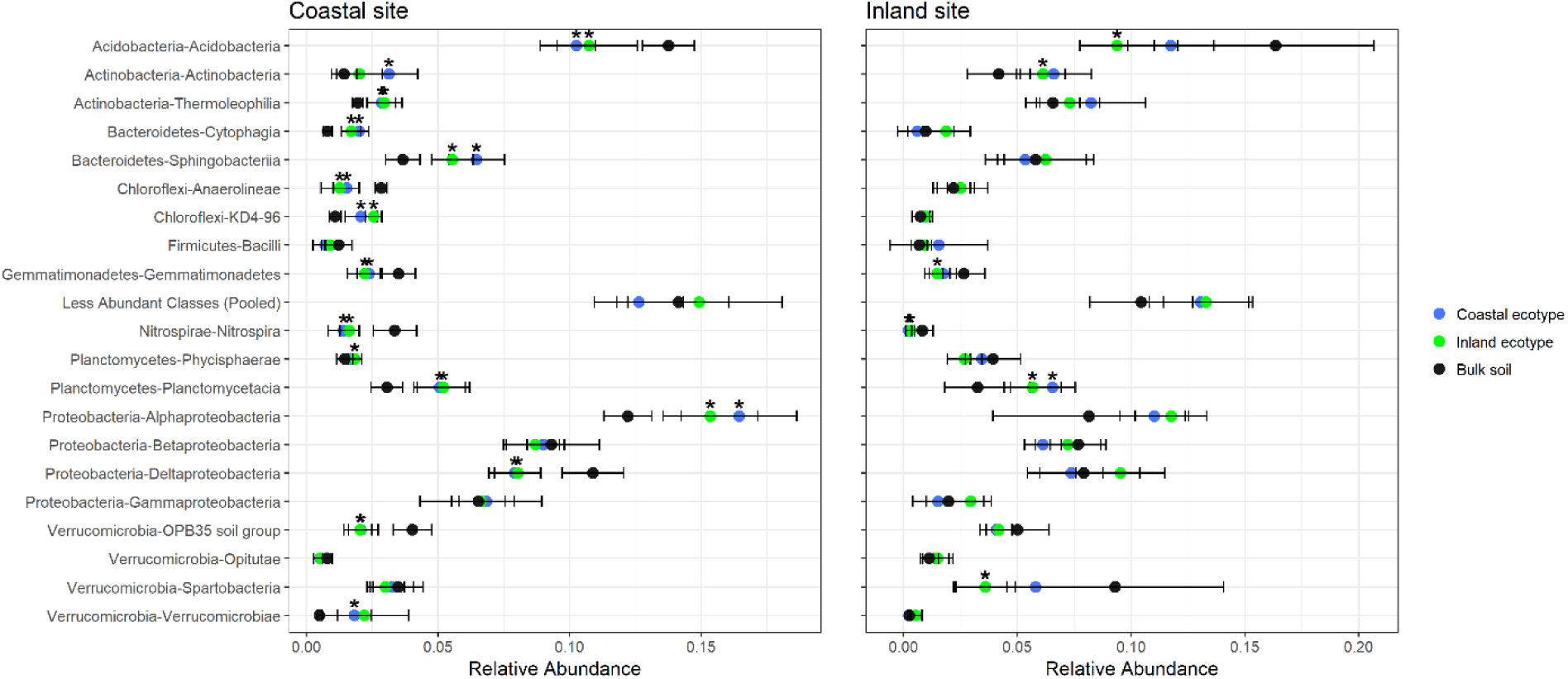
Relative abundance (mean ± SD) of the top 20 most abundant bacterial and archaeal classes in bulk soil and rhizosphere communities of *Mimulus guttatus* planted in two environments. Less abundant taxa were pooled into a single group (“Less Abundant Classes”). Taxa which significantly differed between a specific ecotype and bulk soil are indicated by an asterisk.

Directly comparing the coastal and inland ecotypes (Figure 4), we found that the two ecotypes exhibited very similar relative abundances of microbial taxa at the class level. The two ecotypes did differ in the abundances of several highly abundant taxa, but only at the inland site. At the inland site, the inland ecotype had higher relative abundance of Cytophagia, Deltaproteobacteria, Gammaproteobacteria, and Verrucomicrobiae, but lower relative abundance of Acidobacteria, than the coastal ecotype (Figure 4). Genotypes within each ecotype did not differ in relative abundances of taxa at either the coastal or the inland site (Figure S2).

**Figure 4.**
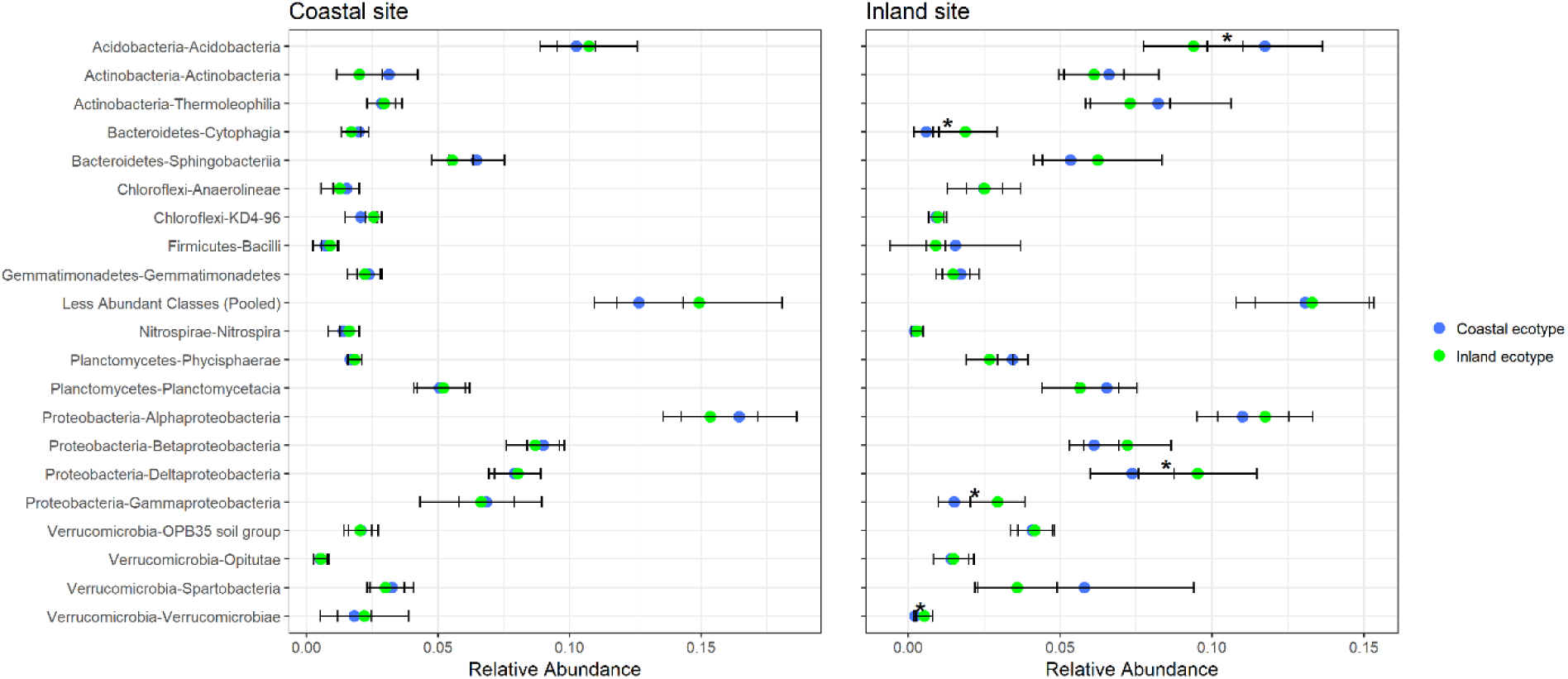
Relative abundance (mean ± SD) of the top 20 most abundant bacterial and archaeal classes in the rhizospheres of coastal (genotypes MRR and SWB pooled) and inland (genotypes LMC and OCC pooled) ecotypes of *Mimulus guttatus* planted in two environments. Less abundant taxa were pooled into a single group (“Less Abundant Classes”). Taxa which significantly differed between ecotypes at a given site are indicated by an asterisk.

### Presence/absence of rare taxa differs between coastal and inland ecotypes at the inland site

Given that inland and coastal ecotypes differed in overall community composition (Figure 1), alpha diversity (Figure 2), and several highly-abundant bacterial classes at the inland site (Figures 3 and 4), but not the coastal site, we further explored the differences between ecotypes at the inland site. Indicator species analysis revealed that no bacterial species were indicative of inland versus coastal ecotypes at the inland site (all adjusted *P*>0.05). In addition, the inland and coastal ecotypes did not differ in relative abundance of any individual OTUs at the inland site. However, the two ecotypes did differ in the presence/absence of numerous OTUs at the inland site: 1,157 OTUs were present in the coastal but not the inland ecotype, while 2,065 OTUs were present in the inland but not the coastal ecotype (Figure 5). These OTUs were in extremely low relative abundance (roughly ten-fold lower mean relative abundance) compared to the 6,290 OTUs shared by the ecotypes and bulk soil. The OTUs distinguishing the coastal and inland ecotypes also had very low occupancy (i.e. were present in a small proportion of samples per ecotype). In the coastal ecotype, only 14 of the 1,157 OTUs unique to the coastal ecotype were present in at least half of the coastal ecotype samples. Similarly, in the inland ecotype, only 99 of the 2,065 OTUs unique to the inland ecotype were present in at least half of the inland ecotype samples. Interestingly, although the majority of the OTUs observed at the inland site (7,537 out of 11,553) were found in bulk soil plus one or both ecotypes, a large number of OTUs were found in either the coastal ecotype (741 OTUs), the inland ecotype (1,234 OTUs), or both (1,484 OTUs), but not the bulk soil. Only 557 of the 11,553 OTUs observed at the inland site were found in bulk soil alone with no observations in either ecotype (Figure 5).

**Figure 5.**
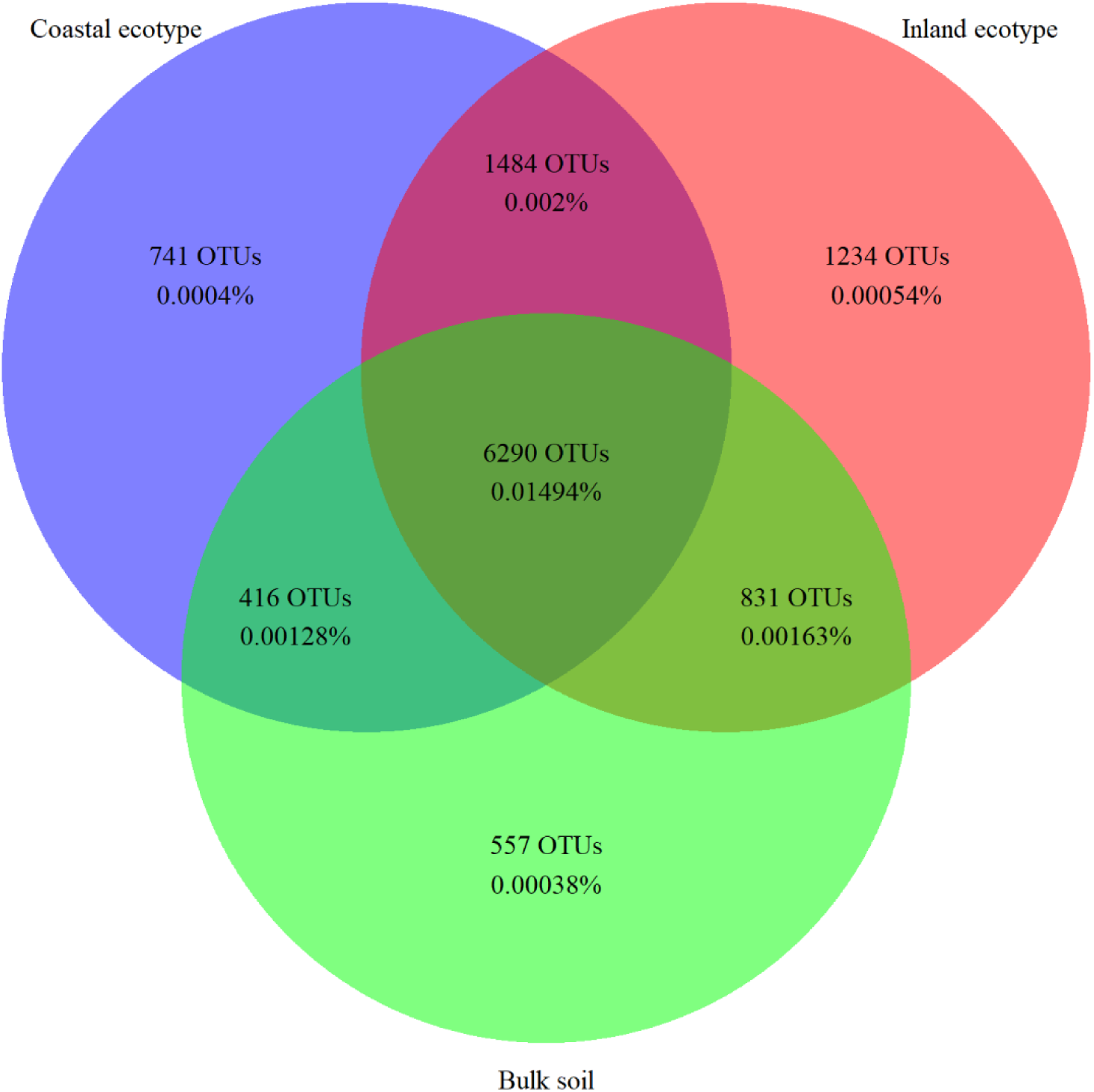
Presence/absence and relative abundance of microbial OTUs in each ecotype rhizosphere and bulk soil at the inland site. Labels indicate the number of OTUs unique to a given set, as well as the mean relative abundance of those OTUs across the full dataset.

## Discussion

Interactions between plant roots and soil microorganisms strongly influence plant health and productivity, yet the relative role of host plant identity versus the local environment in shaping the rhizosphere microbiome is not well understood. To begin to unravel this we examined the rhizosphere communities of two ecotypes of *M. guttatus*, which are locally adapted to distinct environments, in a reciprocal transplant experiment.

The local environment (coastal versus inland site) strongly influenced rhizosphere microbial communities in *M. guttatus*. This effect is due, at least in part, to distinct microbial source pools in the bulk soil at each site. This finding was not surprising given that abiotic conditions strongly differed between the two sites and microbial community structure is often influenced by environmental gradients (Lauber et al. 2009; Fierer et al. 2012; Xue et al. 2018; Sorensen et al. 2019). For example, both salinity (Rath et al. 2019) and moisture availability (Brockett et al. 2012), two of the major factors distinguishing the coastal and inland sites, can have substantial effects on microbial community structure. Nevertheless, despite the drastically different abiotic (soil nutrient availability, salinity, and moisture) and biotic (bulk soil inoculum) conditions between the two sites, the presence of *M. guttatus* strongly influenced microbial communities at both coastal and inland sites. This is in agreement with the general observation that plants play a major role in regulating soil microbial community composition and function (reviewed in (Bulgarelli et al. 2013; Lareen et al. 2016; Coskun et al. 2017).

Host plant identity influenced rhizosphere community composition in *M. guttatus*, but to a smaller extent than the influence of environment. At each site, the two ecotypes exhibited remarkably similar composition of microbial communities at the class level. Many of the shared lineages are commonly associated with rhizospheres, including Actinobacteria, Firmicutes, Alpha- and Beta-proteobacteria (Philippot et al. 2013), suggesting evolutionarily-conserved mechanisms for recruiting and/or sustaining these taxa. Indeed, our results indicate that divergent *M. guttatus* ecotypes recruit phylogenetically similar rhizosphere communities, even in environments to which they are maladapted. Nevertheless, when planted in a common garden at the inland site, the two ecotypes differed in overall community composition, with the inland ecotype recruiting a more OTU-rich and phylogenetically-diverse rhizosphere than the coastal ecotype. This difference in communities between ecotypes at the inland site is largely due to low abundance (rare) and low occupancy (found in a low proportion of samples) microbial OTUs found in one ecotype at the exclusion of the other. Although the relative rarity of these OTUs suggests they may be present in the *M. guttatus* rhizosphere due to stochastic processes rather than by deterministic recruitment by the plant host, rare microbial taxa have the potential to provide a reservoir of microbial functions that can support community stability despite environmental fluctuations (Shade et al. 2014; Shade and Gilbert 2015). The ability of the inland ecotype to harbor greater microbial diversity, due to rare taxa, could potentially contribute to its higher fitness at the inland site compared to the coastal ecotype. Nevertheless, the design of the present study does not allow us to determine whether differing rhizosphere communities at the inland site are a cause or a consequence of the evolutionary divergence between the ecotypes. Future work should explore the potential role of the rhizosphere microbiome in local adaptation in this system by examining growth and fitness of the two ecotypes in sterilized and unsterilized ‘home’ and ‘away’ soil. For this type of experiment, a greater difference in fitness between the two ecotypes in the unsterilized soil would indicate that soil microbial communities contribute to local adaptation and ecotypic divergence in *M. guttatus*.

Taken together, our results indicate that plant host identity impacts rhizosphere communities, and the two locally adapted *M. guttatus* ecotypes are genetically diverged in the factors shaping those communities. Although numerous studies have documented genetic differentiation for rhizosphere microbiome communities in crops and model species in controlled environments (Costa et al. 2006; Micallef et al. 2009; Aira et al. 2010; Peiffer et al. 2013; Mahoney et al. 2017), our work is one of only a few studies reporting genotype-specific effects of wild plants in natural environments (Kuske et al. 2002; Osanai et al. 2013; Aleklett et al. 2015). We hypothesize that variable root exudate composition and/or root morphology between *M. guttatus* ecotypes acts to differentially shape rhizosphere community structure in these ecotypes. Nevertheless, our results show that the effect of host plant identity is environment-dependent, given that the two ecotypes did not differ in community composition when planted at the coastal site. This complex interplay between host identity and environment is in agreement with the contrasting results seen in studies of cultivated crops. For example, some studies report that differences in rhizosphere community composition across species or genotypes are environment-dependent (Marschner et al. 2004; Costa et al. 2006; Peiffer et al. 2013), while others find that differences across species or genotypes are maintained regardless of environment (Mahoney et al. 2017; Marschner et al. 2001). Previous work in the *M. guttatus* system has found that the coastal ecotype exhibits extremely low fitness in inland sites due to near-zero survival-to-flowering rates (Lowry et al. 2008; Lowry and Willis 2010). Although the sample collections made here were completed before the inland site dried out for the summer, it is possible that the early stages of physiological stress at the inland environment contributed to the differences in rhizosphere composition between the two ecotypes seen here. In any case, although the two ecotypes are indeed genetically diverged in factors shaping the rhizosphere microbiome, environmental factors outweigh genetic factors in shaping the *M. guttatus* microbiome at least for the field sites examined in our study.

It is worth noting that numerous taxa were detected in the *M. guttatus* rhizosphere that were not detected in bulk soil. One possible cause of this discrepancy is that the ecotypes recruited taxa that were so rare in the bulk soil that they were below the threshold of detection. Another possibility is that some taxa were carried over from the horticultural soil in which the seedlings were originally germinated before transplanting to the field. A final possibility is maternal packaging of microbial endophytes in the seed (Shade et al. 2017; Rezki et al. 2018), which occurs across diverse plant groups (Nelson 2018) and can influence rhizosphere community composition (Bacilio-Jiménez et al. 2001). More work is needed to determine the potential contributions of seed packaging versus local recruitment to rhizosphere assembly in *Mimulus* and its potential relevance for plant productivity and local adaptation.

In summary, we found that the local environment (coastal versus inland site) strongly influenced rhizosphere communities, at least in part due to distinct composition of the microbial source pool at each site. Although host plant identity also influenced rhizosphere community composition, it was to a much smaller extent than the influence of the environment. At each site, the two ecotypes exhibited remarkably similar composition of microbial communities at the class level, indicating that divergent *M. guttatus* ecotypes recruit phylogenetically similar rhizosphere communities, even in environments to which they are maladapted. Nevertheless, the two ecotypes did differ in rhizosphere community composition at least at the inland site primarily, due to rare (low abundance and low occupancy) OTUs. Overall, the environment-dependence of the differences between ecotypes in rhizosphere communities indicates that strong environmental gradients can obscure plant genetic factors in regulating the *M. guttatus* microbiome. Our findings demonstrate that wild plants strongly impact the structure of soil microbial communities regardless of environment, yet also highlight the context-specific interactions between host identity and local environment in shaping those communities.

## Acknowledgments

The authors thank Benjamin Blackman, Erin Patterson, and the University of California Berkeley greenhouse staff for maintaining our seedlings prior to this experiment and Daniel Jackson for assisting with the fieldwork. We are grateful to the Pepperwood Preserve and University of California, Davis Bodega Marine Reserve for permission to conduct our experiments at these locations. We would especially like to thank Michelle Halbur and Michael Gillogly (Pepperwood) as well as Jacqueline Sones (Bodega) for helping to facilitate our research. The Department of Parks and Recreation of the State of California provided permission to make seed collections for this experiment. This work was supported in part by funding from the Michigan State University Plant Resilience Institute to AS and DBL, and Michigan State University through a startup package to DBL. Computational resources were provided by the Institute for Cyber-Enabled Research.

---

**Supplementary Figure 1.**
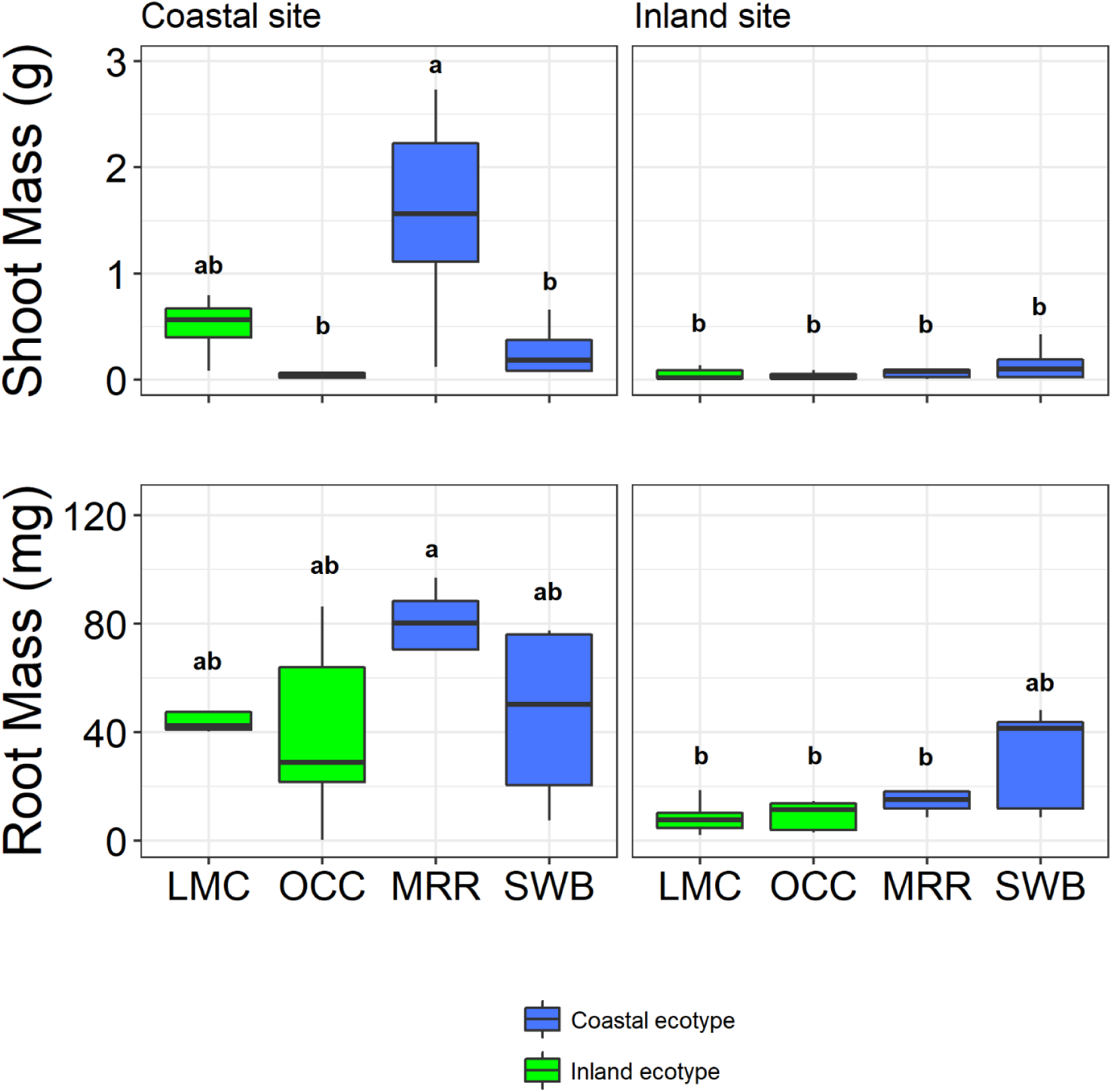
Shoot and root biomass of coastal (MRR, SWB) and inland (LMC, OCC) genotypes of *Mimulus guttatus* planted in two environments. For shoot and root biomass, genotypes that significantly differed are indicated by a different letter above the boxplot.

**Supplementary Figure 2.**
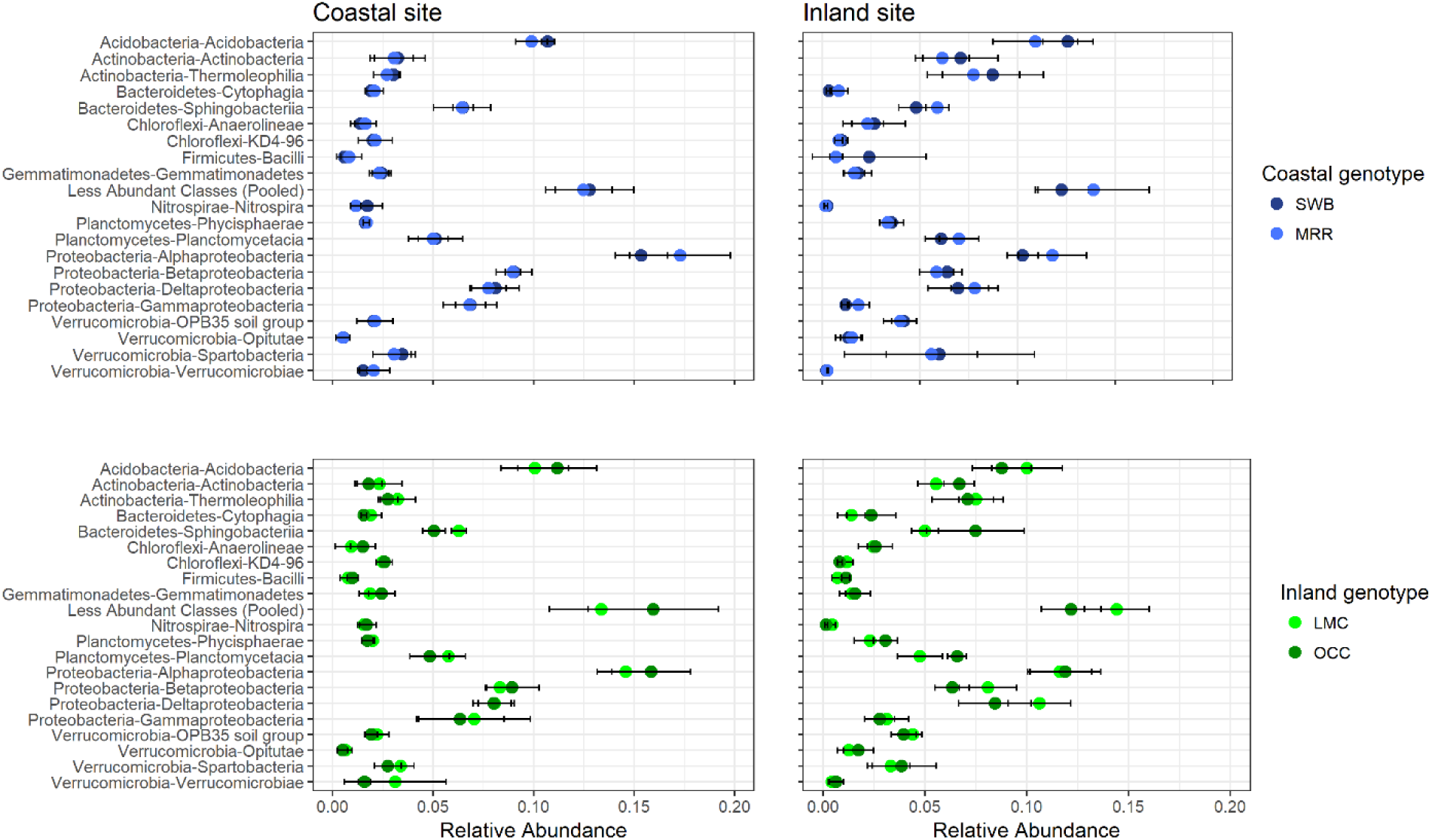
Relative abundance (mean ± SD) of the top 20 most abundant bacterial and archaeal classes in the rhizospheres of coastal (MRR, SWB) and inland (LMC, OCC) genotypes of *Mimulus guttatus* planted in two environments. Less abundant taxa were pooled into a single group (“Less Abundant Classes”). None of the taxa depicted here significantly differed between genotypes at either site.

**Supplementary Table 1.** Soil characteristics (mean ± SE) for bulk and rhizosphere soils collected from *Mimulus guttatus* planted in two environments.

| Soil Variable | Coastal | Inland | p-value |
| --- | --- | --- | --- |
| pH | 6.08 (0.06) | 6.16 (0.07) | 0.3978 |
| Phosphorus (ppm) | 17.4 (1.03) | 3.4 (0.24) | <0.001 |
| Potassium (ppm) | 49.2 (4.14) | 171.4 (7.31) | <0.001 |
| Calcium (ppm) | 788.2 (54.42) | 2518.4 (41.06) | <0.001 |
| Magnesium (ppm) | 227.4 (8.8) | 1603.8 (72.64) | <0.001 |
| Copper (ppm) | 2.42 (0.18) | 21.68 (1.15) | <0.001 |
| Organic Matter (%) | 3.46 (0.26) | 4.9 (0.48) | 0.02891 |
| Sodium (ppm) | 135.8 (7.62) | 50.4 (1.21) | <0.001 |
| Nitrate (ppm) | 0.0 (0.0) | 0.6 (0.23) | 0.05966 |
| Ammonium (ppm) | 5.26 (0.61) | 5.64 (0.69) | 0.6914 |
| Moisture (%) | 34.24 (2.08) | 17.82 (1.82) | <0.001 |
| Total N (%) | 0.1386 (0.02) | 0.1888 (0.02) | 0.1067 |
| Sulfur (ppm) | 23.6 (2.06) | 17.4 (1.29) | 0.03427 |

## Notes

https://github.com/ShadeLab/PAPER_MimulusRecipTransplant_Submitted

